# SPARK: Systematic Profiling of Antibiotic-Phage Response Kinetics

**DOI:** 10.64898/2026.11.06.764789

**Authors:** Brandon A. Berryhill, Teresa Gil-Gil, Dione Sanchez-Hevia, Dan I. Andersson, Nikos Fatsis-Kavalopoulos

## Abstract

Phage therapy is, in practice, combination therapy: phage are almost always added to an ongoing antibiotic regimen, yet the combination is rarely designed to have a specific interaction pattern. Interactions can be antagonistic, additive or synergistic and cannot be inferred from either agent alone. Here we introduce SPARK, a method which quantifies phage–antibiotic interaction from growth curves as the integrated deviation of observed growth from the growth expected under Bliss independence. We profiled 42 combinations spanning the ESKAPE pathogens and *Escherichia coli* with twelve antibiotics. We show that i) synergy and antagonism are resolvable within a single high-throughput assay, with the interaction determined by the phage–antibiotic– bacteria pairing; ii) 21 clinical *Staphylococcus aureus* isolates differ among themselves as widely as the seven species do; and, iii) interactions change between a few hours and two days, several reversing direction. Overall, SPARK needs only a plate reader and the analysis is performed with a specific openly available web tool.

## Introduction

Due to the increasing frequency of infections with antibiotic-resistant bacteria, there has been a resurrection of interest in bacteriophage (phage) for the treatment of bacterial infections ^1, 2^. In 2019, resistant infections were directly responsible for an estimated 1.27 million deaths worldwide ^3^, and the pipeline of new antibiotics has not kept pace with the spread of resistance ^4,5^. Phages are specific to their hosts, replicate at the site of infection, and kill by mechanisms unrelated to those of antibiotics, which makes them attractive for infections that no longer respond to conventional treatment ^1, 2^. In current practice, however, phages are almost never given alone; in the majority of reported compassionate-use cases they are administered on top of an ongoing antibiotic regimen ^2, 6, 7^. Phage therapy is thus, in practice, combination therapy but the combination is rarely rationally designed ^6, 7^. Clinicians add phage as an empirical adjuvant to the regimen the patient is already receiving, and the partner antibiotic(s), the dose, and the timing are seldom informed by any measurement of how the two agents act together against the infecting strain ^1, 6, 8^.

The joint action of phage and antibiotics can be expected to show both synergies and antagonisms. Sub-inhibitory concentrations of some antibiotics have been shown to markedly enhance phage replication and bacterial killing, a phenomenon termed phage–antibiotic synergy ^9–12^; whereas others, particularly those that inhibit protein synthesis, suppress phage amplification, and reduce the combined effect below that of either agent alone ^13, 14^. Synergy and antagonism can arise with the same phage depending on the antibiotic and with the same antibiotic depending on the host ^14–16^. The outcome of a given combination therefore cannot be inferred from the activity of each agent separately, rather it must be measured. There is currently no method that quantifies these interactions in a way that captures effects in both directions, describes how interactions change over the course of treatment, and scales to the number of strains, phage, and antibiotics relevant to a clinical decision ^8, 14, 17^. Consequently, there is no rational basis for deciding which phage and antibiotics to combine.

The difficulty is in part conceptual, as the methods used to measure antibiotic–antibiotic interactions do not transfer to a self-amplifying agent. Checkerboard assays and the fractional inhibitory concentration index derived from them depend on the minimum inhibitory concentration (MIC) of each agent ^18, 19^. A phage has no MIC in the conventional sense: it replicates on its host, so its effective dose changes during the experiment and depends on the density and physiological state of the bacterial population ^14, 20^. A single read-out of the interaction at 16 to 24 h is therefore an arbitrary point in a dynamic process. Time–kill assays capture these dynamics directly but rely on colony counts (something which does not translate to the clinical lab) and are difficult to scale beyond a small number of combinations. Plaque-based assays, including those from which phage–antibiotic synergy was first described, report on phage propagation rather than on the fate of the bacterial population ^9, 11^, and matrix-based approaches such as synograms extend the checkerboard design to phage titres while remaining tied to a single endpoint ^14^. The same constraint applies to the reference model used to calculate these interactions. Loewe additivity, which underlies isobologram and fractional inhibitory concentration analyses, defines non-interaction through dose equivalence ^21^, and dose equivalence has no counterpart for an agent whose effective dose is not fixed. Bliss independence instead derives the expected combined effect from the fractional effect of each agent measured alone ^21, 22^. We have previously applied this measurement in the sublethal interaction factor (SIF) which applies Bliss independence to complete growth curves to quantify antibiotic–antibiotic interactions ^23^.

Here we employ growth curves and Bliss independence to measure the joint action of phage and antibiotics, in an approach we call SPARK (Systematic Profiling of Antibiotic–Phage Response Kinetics). Growth under combined treatment is compared at every timepoint with the growth expected of the two agents acting independently and the difference is integrated over the experiment to give the value of the SIF for that combination. We restrict our consideration to sub-inhibitory concentrations of the antibiotic at which both agents act on a growing population. We demonstrate that: i) the method resolves synergy and antagonism across 42 combinations spanning the six ESKAPE pathogens (*Enterococcus faecium, Staphylococcus aureus, Klebsiella pneumoniae, Acinetobacter baumannii, Pseudomonas aeruginosa, and Enterobacte*r species) ^3, 24^ and *Escherichia coli* with twelve antibiotics; ii) the same antibiotic produces synergy in one species and antagonism in another, so that an interaction measured in one host does not transfer to the next; iii) interaction outcomes differ among 21 clinical isolates of *Staphylococcus aureus* as much as they differ between species; and, iv) interactions change over time from a few hours to nearly two days after treatment and in several cases reverse direction during the experiment, behaviors which would not have been seen via endpoint measurements. Our approach requires only a plate reader using an assay similar to those already run in most laboratories. This method can be employed for any phage and bacterium that can be cultured in vitro.

## Results

### SPARK quantifies phage–antibiotic interaction from complete growth curves

SPARK compares the growth of a bacterial population under combined phage and antibiotic treatment with the growth expected if the two agents acted independently. Each combination is profiled under four conditions on the same microtiter plate, untreated control, phage alone, antibiotic alone, and the combination, with optical density read continuously over 48 h (Fig. 1a and 1b). Antibiotics are used at sub-inhibitory concentrations, which produce measurable but incomplete inhibition and leave a growing population on which the phage can act, and phage is added at a fixed multiplicity of infection. Each condition is normalized to the untreated control at every time point, and the expectation of independent action is constructed from the two single-agent curves of the same experiment (Fig. 1c and Online Methods).

**Figure 1.**
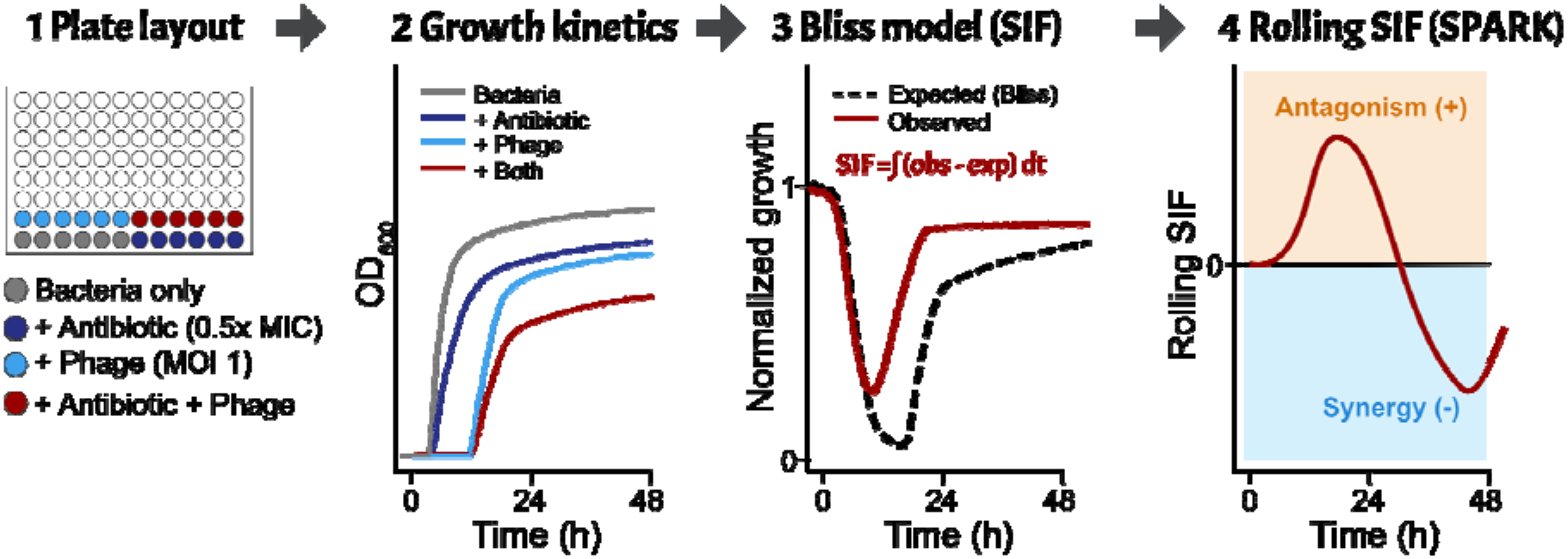
The SPARK assay and the interaction metric. (a,b) Each phage–antibiotic combination is profiled in four conditions on one plate: untreated control, phage alone, antibiotic alone, and the combination, with optical density recorded continuously over 48 h. Each condition is normalized to the untreated control at every time point, and the expectation of independent action is the product of the two single-agent curves. (c) The interaction score is the integrated difference between the observed and expected curves, ΔAUC = AUC(experimental) − AUC(theoretical), read either as a single endpoint value or, accumulated from the start of the experiment to each time point, as a trajectory. (d) Blue, less growth than expected (synergy); red, more growth than expected (antagonism); grey, the neutral interval.

Interaction strength is the integrated deviation of the observed combination from that expectation. The difference in area under the growth curve, ΔAUC = AUC(experimental) − AUC(theoretical), is the value of the sublethal interaction factor (SIF) for that combination: negative values indicate synergy, positive values antagonism, and values within the neutral interval are treated as independent action. Because the same four curves also support a cumulative calculation, one experiment yields either a single endpoint score or a trajectory of interaction over time (Fig. 1d). Together, this design gives a measurement that requires only a plate reader and that is internally controlled for the isolate, the phage, and the antibiotic concentration used.

### SPARK resolves antibiotic- and species-specific interactions across the ESKAPE pathogens and *E. coli*

To establish the measurement range of the method and to test whether it recovers interactions of both directions, we profiled 42 phage–antibiotic combinations spanning the six ESKAPE pathogens and *Escherichia coli* (Online Methods). Shown in Fig. 2a are the mean endpoint values for all 42, each species receiving four drugs which were given to all seven along with two further agents selected for clinical relevance to that organism. Phages which were previously used for phage therapy were preferred, see Extended Data Table 1 for more information.

**Figure 2.**
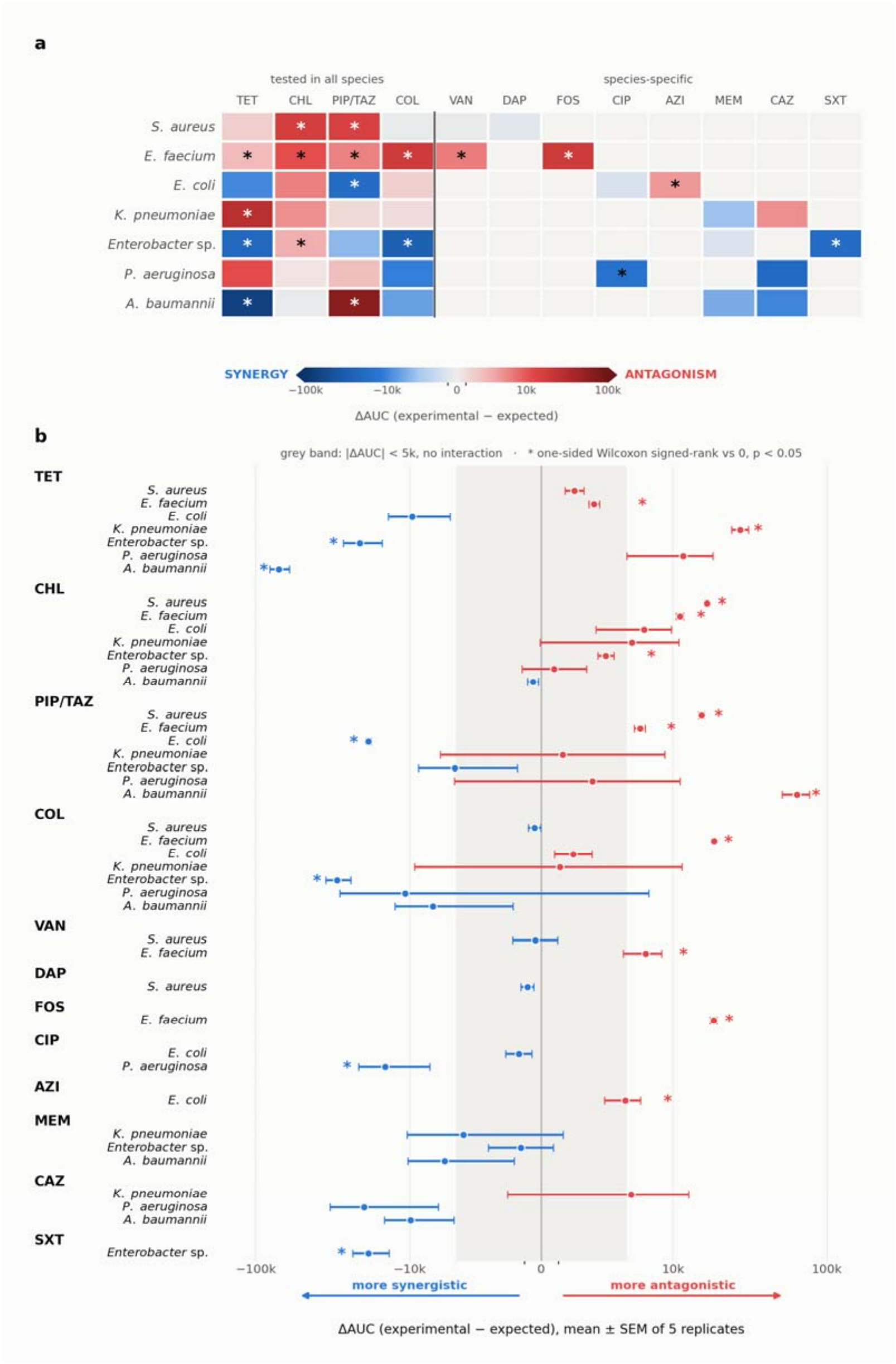
Phage–antibiotic interactions across the ESKAPE pathogens and *Escherichia coli*. (a) Mean ΔAUC for 42 phage–antibiotic combinations, seven species at six antibiotics each. Rows are species and columns antibiotics, with TET, CHL, TZP, and COL, given to every species, left of the divider, and the species-specific agents to the right: VAN and FOS for E. faecium; VAN and DAP for *S. aureus*; CAZ and MEM for *K. pneumoniae* and for A. baumannii; CIP and CAZ for P. aeruginosa; MEM and SXT for Enterobacter sp.; CIP and AZI for E. coli. Cells outside this design were not tested. (b) The same values as mean ± s.e.m., grouped by antibiotic. Grey band, |ΔAUC| < 5,000. Blue, less growth than expected (synergy); fred, more growth than expected (antagonism). n = 5 independent replicates; asterisks mark ΔAUC differing from zero (one-sided Wilcoxon signed-rank test, p < 0.05), 18 of 42 combinations. See Online Methods for the metric, the neutral interval, and the axis scaling.

The assay returned interaction values spanning more than three orders of magnitude (Fig. 2b), from −70,789 for *A. baumannii* with tetracycline (TET) to +63,996 for the same species with piperacillin–tazobactam (TZP). The fact that half of the interactions were neutral or non-interacting we take as evidence that the metric returns values centered on zero when no interaction is present, rather than defaulting to a direction.

Reading down the columns of Fig. 2a, we find that a single antibiotic can produce opposite interactions in different bacterial backgrounds. TET divided the panel evenly, with strong synergy in *A. baumannii* (−70,789, p < 0.05) and *Enterobacter* sp. (−21,042, p < 0.05) and antagonism in *E. faecium* (+3,082, p < 0.05) and *P. aeruginosa* (+11,643). TZP was similarly divided, being the strongest antagonistic combination in *A. baumannii* (+63,996, p < 0.05) while producing significant synergy in *E. coli* (−18,503, p < 0.05). Colistin (COL) was synergistic in *Enterobacter* sp. (−29,655, p < 0.05) but the most antagonistic combination tested in *E. faecium* (+18,481, p < 0.05). Chloramphenicol (CHL) was the only antibiotic with a consistent direction, showing antagonism in all cases.

Reading along the rows, we also find systematic differences between bacterial backgrounds. *E. faecium* was antagonistic in all six combinations tested, all of which reached significance, and was the only species in the panel with a uniform interaction direction. *A. baumannii* spanned the full measurement range, containing both the most synergistic and the most antagonistic combination in the study. *P. aeruginosa* returned only one significant interaction, with ciprofloxacin (CIP; −14,480, p < 0.05); its remaining combinations were not distinguishable from independence, with COL spanning −63,518 to +41,897 across five replicates in this bacterium. We postulate that this dispersion reflects variation between replicate cultures rather than a small underlying effect. We discuss these consequences for the detection limit of our assay below.

Taken together, these data show that interaction outcome is a joint property of the antibiotic and the bacterial background, so that neither agent alone predicts it.

### Rolling integration resolves when an interaction emerges and whether it reverses

A single endpoint measurement collapses an inherently dynamic process into one number. The four curves of each experiment already contain continuous time course data. We therefore applied the rolling formulation of SIF to the same 42 combinations, expressing the difference between the cumulative observed and expected AUC at each time point as a percentage of expected (Online Methods). Shown in Fig. 3a are the resulting trajectories.

**Figure 3.**
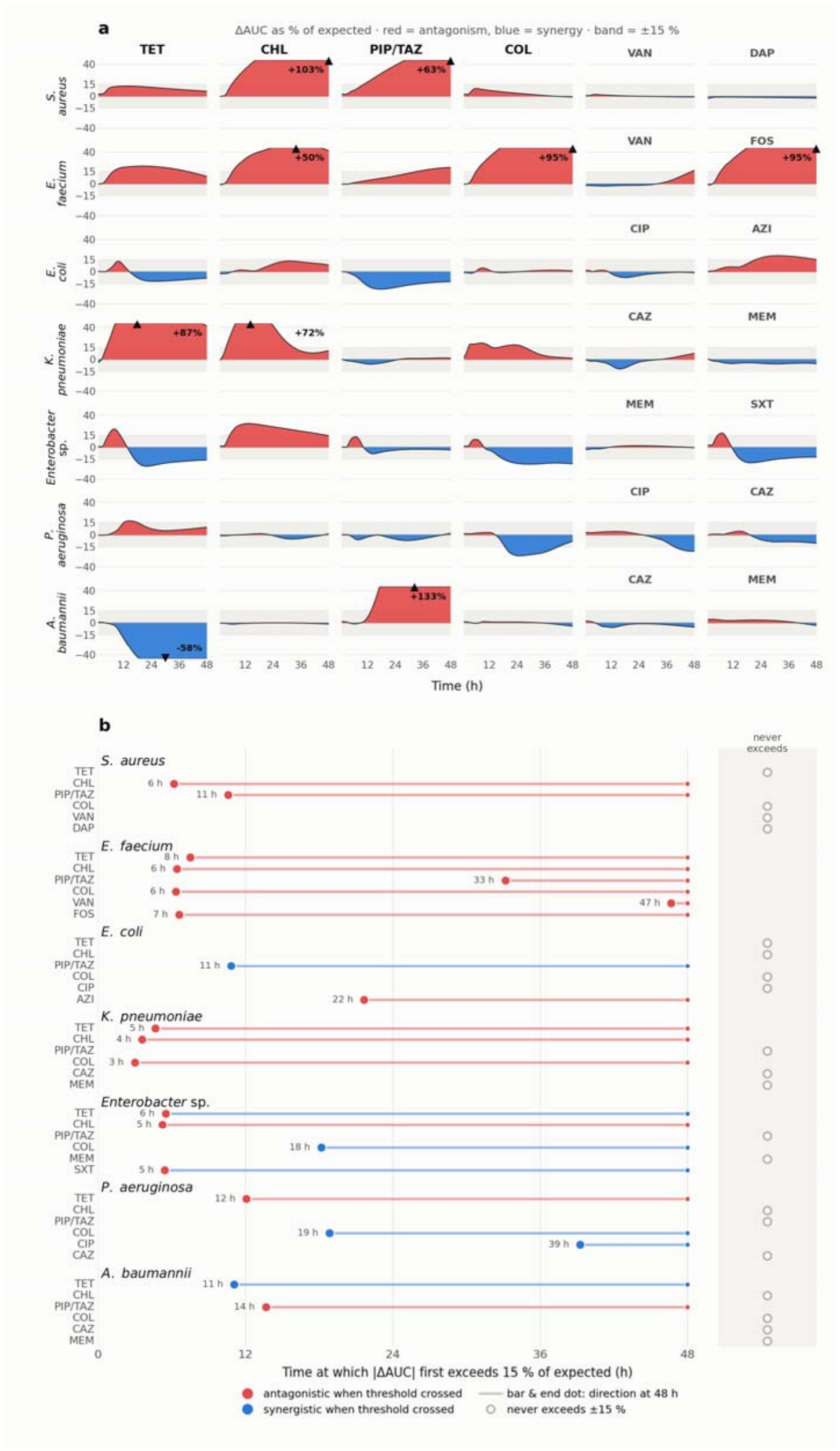
Time course of phage–antibiotic interaction across seven pathogens and twelve antibiotics. (a) Rolling ΔAUC, as a percentage of expected, over 48 h for all 42 combinations; rows are species, the first four columns TET, CHL, TZP, and COL, given to every species, and the last two that species’ own agents as listed for Fig. 2. The y-axis is truncated at ±45 %; triangles mark the 9 combinations exceeding it, with their maximum deviation. (b) Time at which each combination first exceeds ±15 % of expected for at least 1 h (large dot) and its direction at 48 h (terminal dot); open symbols, combinations within ±15 % throughout. Grey band, ±15 %. Blue, synergy; red, antagonism. n = 5 independent replicates.

We found that the rolling metric converged on the same classifications as the endpoint measurement while adding temporal structure. Combinations identified as strongly interacting in the endpoint panel were recovered in the time-resolved data with matching direction, including synergy for *A. baumannii* with TET (−52 % of expected at 48 h) and for *E. coli* with TZP (−12%), and antagonism for *A. baumannii* with TZP (+95 %) and across all *E. faecium* combinations. The principal information gained by time resolution was the timing of divergence, which varied by an order of magnitude across the panel. Shown in Fig. 3b is the time at which each combination first left the neutral band: 22 of the 42 did so at some point during the experiment and 20 remained within it throughout. Among those that diverged, the earliest did so within the first six hours, *Enterobacter* sp. with trimethoprim–sulfamethoxazole (SXT) at 5.4 h, *Enterobacter* sp. with CHL at 5.3 h and *E. faecium* with CHL at 6.4 h, whereas others moved out of the neutral band only in the final third of the experiment, including *P. aeruginosa* with CIP at 39 h and *E. faecium* with vancomycin (VAN) at 47 h. We therefore take the time of divergence to be a property of the combination in its own right. A measurement at a fixed early time point would have classified a substantial fraction of these combinations as non-interacting.

Time resolution additionally revealed trajectories that are not monotonic and whose direction changes during the experiment, a behavior that cannot be represented by any single endpoint value. *Enterobacter* sp. with TET rose to +22.7 % at 7.9 h before falling to −16.0 % by 48 h, and the same species with SXT followed a comparable path (+17.8 % at 7.1 h to −12.1 %). *E. coli* with TET showed the same pattern at smaller amplitude (+13.3 % at 9.8 h to −7.9 %), while *P. aeruginosa* with COL diverged to −24.1 % at 24.3 h before returning to within the neutral band by the end of the assay. In contrast, several combinations diverged progressively and without reversal, most prominently *A. baumannii* with TZP, which rose steadily from 13.6 h to a maximum of +133 % at 32.8 h (Fig. 3a). We interpret the reversing trajectories as evidence that the process dominating growth of the combined culture changes during the assay, so that the direction of an interaction is not fixed at the point it first becomes detectable.

Overall, the rolling formulation reproduces the endpoint classifications while additionally reporting when an interaction emerges and whether its direction is stable.

### Within-species profiling resolves isolate-level variation in interaction outcome

The panel above uses one representative isolate per pathogen, as most published comparisons of phage–antibiotic combinations do. To determine whether a species-level measurement is representative of the species, we applied the endpoint metric to 21 clinical *S. aureus* isolates, each tested with phage in combination with vancomycin and with piperacillin–tazobactam. Shown in Fig. 4a are the mean values for all 42 isolate–antibiotic combinations.

**Figure 4.**
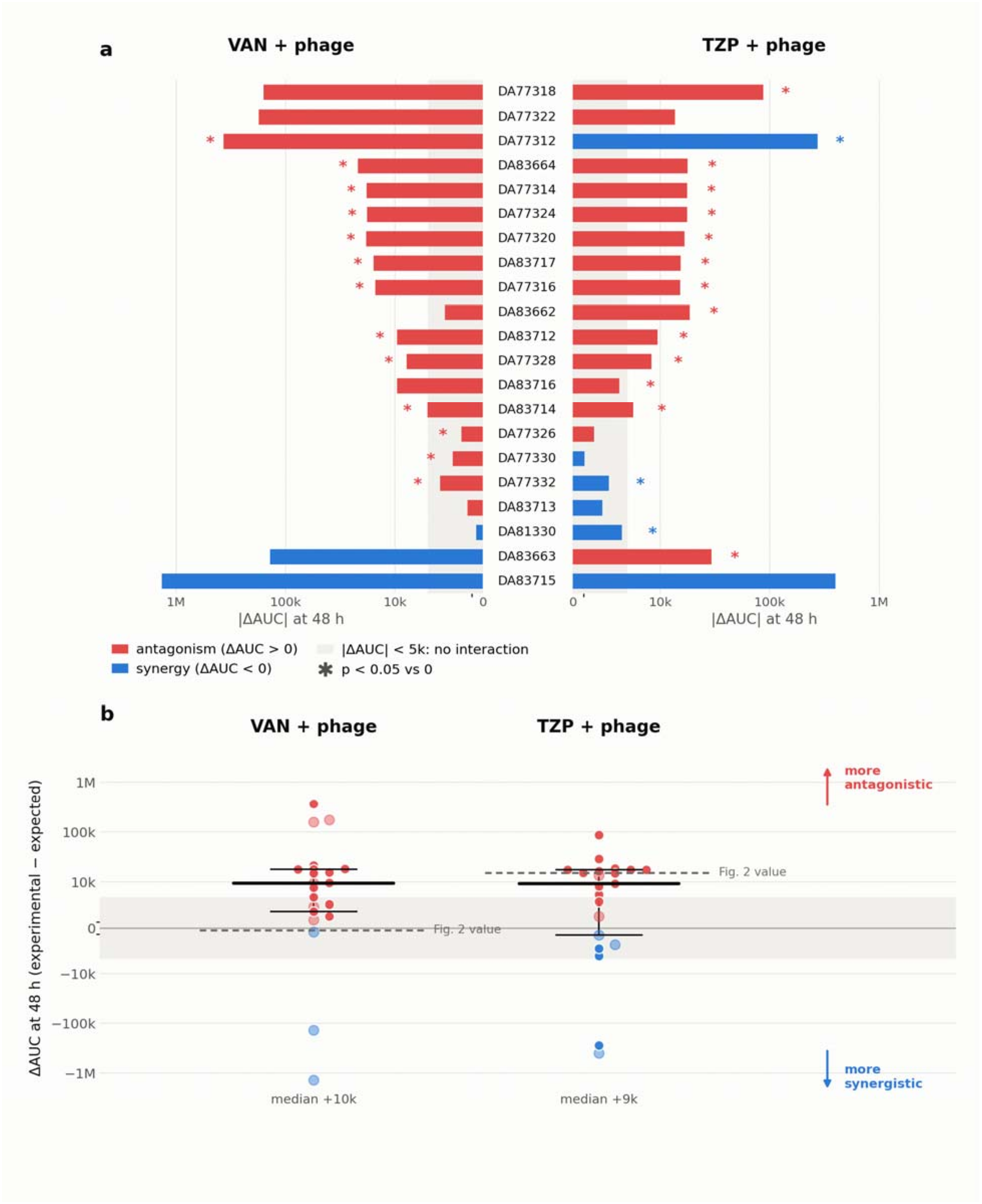
Strain-to-strain variation in phage–antibiotic interaction across 21 *Staphylococcus aureus* clinical isolates. (a) Mean ΔAUC per isolate for phage with vancomycin (VAN, left of the labels) and piperacillin–tazobactam (TZP, right), ordered from most antagonistic to most synergistic; bars of differing color mark an isolate that reverses direction between the two antibiotics. (b) Distribution of the same isolate means, with median and interquartile range; filled points are significant, and the dashed line gives the species-level value for *S. aureus* from Fig. 2. Blue, synergy; red, antagonism. n = 5 independent replicates; asterisks, one-sided Wilcoxon signed-rank test, p < 0.05 (13 of 21 isolates with VAN, 16 of 21 with TZP).

Interaction values within this single species spanned a range comparable to that observed across the entire seven-species panel (Fig. 4b), from −1,356,609 (DA83715 with VAN) to +367,953 (DA77312 with VAN). Twenty-nine of the 42 combinations differed significantly from independent action, 13 of 21 isolates with VAN and 16 of 21 with TZP. Antagonism predominated: 18 of 21 isolates were antagonistic with VAN and 15 of 21 with TZP, with median values of +10,000 and +9,000 respectively.

We find that isolate identity, rather than the antibiotic, was the principal determinant of outcome (Fig. 4a). Fourteen isolates were antagonistic with both antibiotics and two were synergistic with both, while five isolates reversed direction between the two drugs. The most pronounced of these was DA77312 (Fig. 4a), which was significantly antagonistic with VAN (+367,953) and significantly synergistic with TZP (−272,831); DA83663 showed the opposite pattern, trending synergistic with VAN (−137,895) and significantly antagonistic with TZP (+29,172). Interaction values for the two antibiotics were only moderately associated across isolates (Pearson r = 0.64, Spearman ρ = 0.39), indicating that the rank order of isolates was not preserved between drugs.

Comparison with the species-level measurement demonstrates the consequence of profiling a single isolate. The *S. aureus* values obtained in the seven-species panel (VAN, −364, not significant; TZP, +15,335, p < 0.05) sit close to the isolate medians reported here, yet the VAN measurement alone would have classified the species as non-interacting, whereas 13 of 21 individual isolates showed statistically significant antagonism, and two isolates showed synergy exceeding 100,000 in magnitude.

Replicate dispersion was concentrated among the largest-magnitude isolates. DA83715, the most synergistic isolate in the panel, had a standard error of 815,448 with VAN and did not reach significance, and DA77322 returned p = 0.50 because four replicates lay near zero while a fifth reached +936,704. Conversely, isolates in the +10,000 to +20,000 range were highly reproducible, with standard errors below 1,000 in several cases. We take this to mean that effects above roughly 10^5^ in absolute value require additional replication before they can be classified with confidence, and that the detection limit of the endpoint metric is set by replicate variance rather than by the size of the effect.

Collectively, these data establish that a single isolate is not a reliable proxy for the interaction behavior of its species.

### Time-resolved profiling within a species reveals isolate-specific interaction kinetics

Finally, we asked whether the temporal behavior seen across species is also present within one. We applied the rolling formulation to the same isolate panel, generating normalized interaction trajectories for all 42 isolate–antibiotic combinations. Shown in Fig. 5a, c are the isolates whose deviation changes during the time course, and in Fig. 5b, d the remainder.

**Figure 5.**
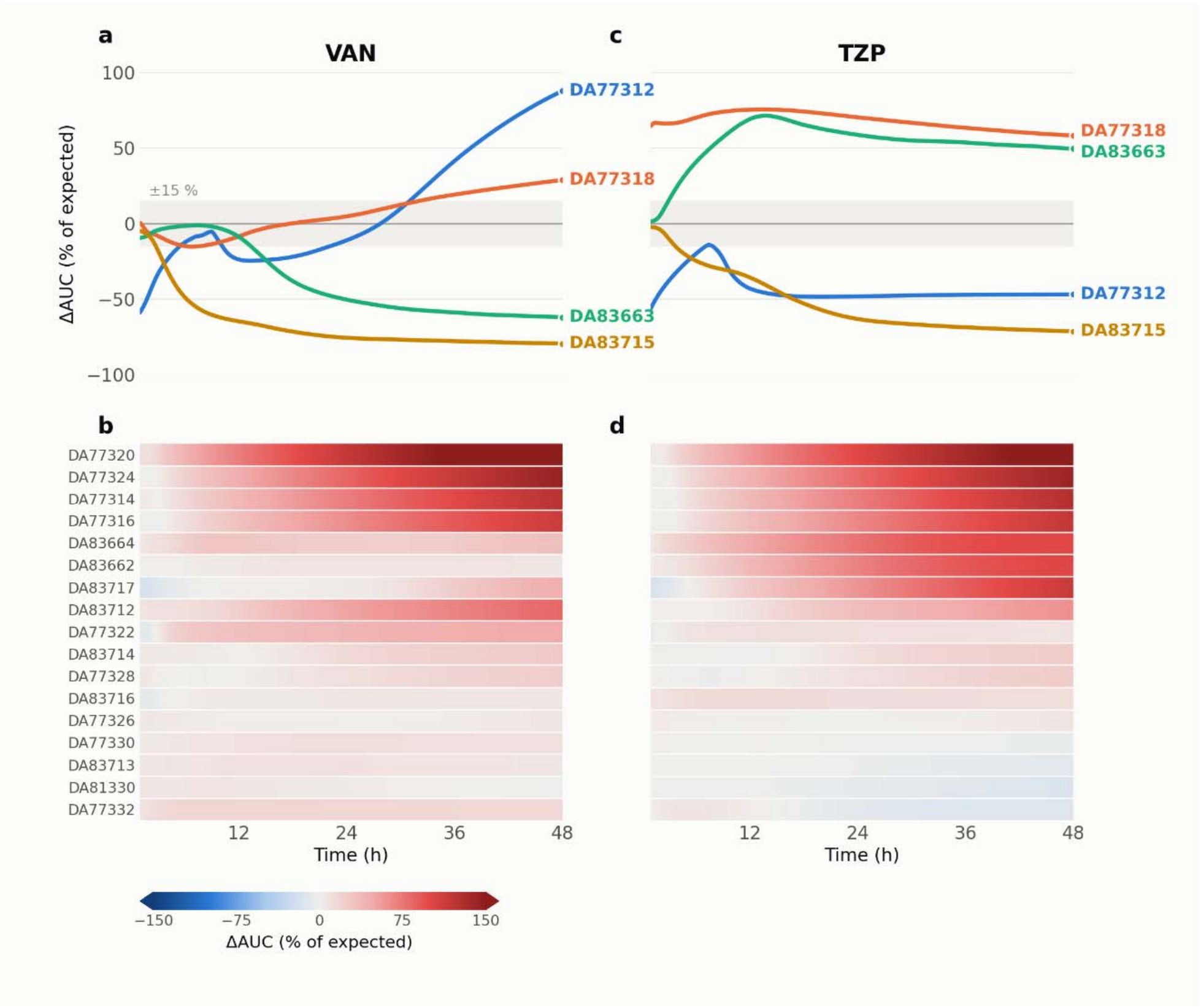
Time-resolved deviation from expected growth for phage–antibiotic combinations across 21 *S. aureus* clinical isolates. Rolling ΔAUC, as a percentage of expected, over 48 h. (a, c) Isolates whose deviation changes direction or magnitude during the time course, for the vancomycin (VAN, a) and piperacillin–tazobactam (TZP, c) combinations; points mark the 48 h endpoint and the grey band ±15 %. (b, d) The remaining isolates for the same two combinations, with color giving the same quantity (scale capped at ±150 %); row order is the clustering of each isolate’s joint VAN and TZP trajectory and is the same in both panels. Isolates are identified by DA number. Blue, synergy; red, antagonism. n = 5 independent replicates.

We found the endpoint and time-resolved analyses mutually consistent: the direction of the normalized value at 48 h agreed with the sign of the endpoint ΔAUC for all 21 isolates with both antibiotics, and the two measures were strongly rank-correlated (Spearman ρ = 0.82 for VAN and 0.83 for TZP). Thirty of the 42 combinations left the ±15 % neutral band during the experiment, while 12 remained within it throughout (Fig. 5b, d).

As in the multi-species panel, we found that the time at which an interaction became detectable varied widely between isolates challenged with the same antibiotic. Divergence times ranged from under three hours (DA83664 and DA77320 with TZP, 2.6 and 2.8 h) to beyond twenty hours (DA83714 with VAN at 23.9 h, DA77328 with TZP at 30.6 h), a span of more than an order of magnitude within a single species and a single drug pairing. Four isolates crossed the neutral band before three hours, whereas four others required more than twenty. We read this spread as the within-species counterpart of the divergence times measured across species, and it is of the same order.

We found the magnitude of antagonism similarly isolate-dependent, and in several cases substantially larger than any value observed in the multi-species panel: DA77320 reached +191% of expected with VAN and +169 % with TZP, followed by DA77324 (+145 % and +143 %), DA77314 (+126 % and +129 %) and DA77316 (+115 % and +119 %). At the opposite extreme, DA83715 diverged early and synergistically with both antibiotics, reaching −49 % of expected by 6 h with VAN and ending at −80 %, and −71 % with TZP (Fig. 5a, c).

Three isolates reversed the direction of their interaction during the experiment (Fig. 5a, c). DA77312 remained below expectation for the first thirty hours with VAN, crossing zero at approximately 30 h and rising to +88 % by 48 h, while with TZP the same isolate fell to −45 % within 12 h and remained synergistic for the rest of the assay. DA83717 and DA77318 followed similar reversing trajectories with VAN, ending at +46 % and +29 % of expected after early synergistic phases. DA83663 illustrates the converse case of a stable but late interaction, tracking the expected curve for the first twelve hours with VAN before falling to −62 %, while with TZP it rose to +71 % by 14 h and settled at +49 %.

Together, these results show that both the timing and the direction of an interaction are properties of the individual isolate rather than of the species.

## Discussion

Critical to the rational design of phage–antibiotic combination therapy is the measurement of how the two agents act together against the infecting strain. Here we have employed complete growth curves and Bliss independence to provide a method to generate this measurement. In an approach we call SPARK, we have shown that: i) a single integrated score resolves synergy and antagonism across 42 combinations of phage and antibiotics spanning the ESKAPE pathogens and *Escherichia coli*; ii) direction is set by the pairing rather than by the drug; iii) 21 clinical isolates of *S. aureus* differ among themselves as widely as the seven species differ from one another; and, iv) interactions appear between a few hours and nearly two days after treatment, and several reverse direction before it ends.

Several properties of the method underpin its practical value. First, SPARK requires only a temperature-controlled plate reader and a multi-well plate, so that panels of clinical isolates rather than single strains become tractable. Second, one experiment yields both an endpoint score and a full trajectory from the same curves, so the read-out time need not be chosen in advance. Third, the expectation is built at every time point from the single-agent curves of the same plate, so each measurement is internally controlled and no inhibitory-concentration parameter is required for the phage ^14, 23^. Fourth, the score integrates the whole trajectory, so that lag modulation, growth inhibition, lysis, and regrowth each contribute to one quantity. The analysis itself runs as a web tool that takes raw plate-reader output and returns both the endpoint score and the trajectory, so neither scripting nor local installation is required.

The most consequential observation the method enables is that phage–antibiotic interaction is a time-dependent property rather than a fixed one. *Enterobacter* sp. with tetracycline was antagonistic early, reaching +22.7 % of expected growth at 7.9 h, before becoming synergistic by 48 h; *P. aeruginosa* with colistin diverged to −24.1 % at 24 h and then returned within the neutral interval. Detection times ranged from approximately 5 to 47 h across species, and from under 3 to more than 30 h among isolates of one species given the same antibiotic. Two consequences follow. First, the classification of a combination depends on when it is observed: a read-out at 6 to 8 h would have missed most of the interactions detected here. Second, opposing phases can cancel within the integrated endpoint, so a value close to zero does not by itself indicate independent action. Each inflection marks a change in the process dominating growth and is informative in its own right.

We focused the within-species analysis on *S. aureus* because it is among the most frequent targets of clinical phage therapy ^6, 13^ and because well-characterized isolate collections are available ^25, 26^. The range of interaction values within this single species was as wide as that across all seven. Five isolates reversed direction between the two antibiotics, the rank order of isolates was only weakly preserved between drugs, and the species-level vancomycin measurement was indistinguishable from independent action while 13 of 21 isolates were significantly antagonistic. We interpret this to mean that phage–antibiotic interaction is a property of the individual isolate, much as antimicrobial susceptibility is, rather than of the species ^16, 27^. Claims of synergy or antagonism drawn from a single strain should accordingly be treated with caution ^17, 25^.

These findings bear on how phage therapy is delivered. Phages are rarely administered alone; in most reported cases they are given on top of ongoing antibiotic therapy ^6, 8^. The practical question is therefore which antibiotic the patient is being treated with when the phage is brought onboard. Antagonism predominated, accounting for 12of 18 significant interactions in the species panel and 18 of 21 *S. aureus* isolates given vancomycin. Should this obtain in vivo, a concurrently administered antibiotic could reduce or nullify the contribution of the phage, and timing may matter as much as the choice of partner as seen by the variation in interaction over time. We accordingly recommend that the partner antibiotic be chosen against the patient’s own isolate and therapeutic phage, which the format of SPARK makes feasible prior to the start of treatment.

Although this study was not designed to resolve mechanism, we postulate that these dynamics follow from known features of phage–host biology. Phage amplification depends on bacterial physiology, with its growth rate influencing latent period and burst size ^11, 28, 29^, so antibiotics that impair translation would be expected to reduce phage productivity, consistent with the antagonism we observed for chloramphenicol. Drug class was nevertheless unpredictive on its own: tetracycline was strongly synergistic in *A. baumannii* and *Enterobacter* sp. yet antagonistic in *E. faecium* and *P. aeruginosa*. Late inflections occurred on the time scale over which phage-resistant subpopulations re-emerge, which could explain the late divergences ^13, 30, 31^. β-lactam-induced filamentation has been proposed as a route to enhanced phage production ^9, 10^, and piperacillin–tazobactam was antagonistic in three species yet synergistic in *E. coli*. Possibly degree ofhost filamentation at the concentration used determines the direction of a β-lactam interaction. Testing this will require tracking phage titres and resistance through the assay, at the time points the trajectories identify.

We recognize some limitations to our study. Each combination was tested at one sub-inhibitory concentration, one multiplicity of infection, and one inoculum density. Both the magnitude and direction of an interaction may depend on all three. Each species was paired with a single phage, so the values describe specific phage–antibiotic-host combinations rather than properties of a drug or a species. The thresholds are themselves inferred: the neutral interval, and the direction calls that follow from it, were set against replicate variance rather than against any clinical outcome, so synergy and antagonism here are statements about growth in vitro and not about treatment success. Optical density reports biomass rather than viable cell number, so filamentation, lysis debris, and aggregation all affect the read-out ^32^. Given these limitations, we do not believe these limits require a different assay; each is a matter of experimental scope and limits the interpretation of the data we generated for the validation of the assay rather than of the method itself.

The Bliss independence calculation treats the two agents symmetrically, so the score gives the direction and magnitude of an interaction but does not determine which agent drives it. Resolving that directionality is the natural next step, which could be done in two ways: first, profiling small matrices of concentration and multiplicity of infection, from which agent-specific potency shifts can be derived ^14^; and second, analyzing which single-agent trajectory the combination follows at each inflection. We also intend to compare sequential with simultaneous administration of both agents, which have been described to alter treatment outcome for *S. aureus* ^13, 33^, and to adapt the read-out to biofilm formats, where the clinical case for combination therapy is strongest ^11, 33, 34^. The most critical (and largest undertaking) next step is to attach clinical meaning to the readout values. Phage is increasingly delivered under compassionate-use protocols that record the isolate, the phage, the partner antibiotic, and the outcome ^2, 6^; profiling those combinations at scale would allow SIF breakpoints to be derived as MIC breakpoints were, from the distribution of values against observed outcome rather than against replicate variance.

## Online Methods

### Bacterial strains and bacteriophage

The seven species of the panel and their phage are listed in Extended Data Table 1. The bacterial strains are the ATCC type strain of each species, obtained from ATCC; the phage and their sources are given in the same table. The within-species panel comprised 21 clinical S. aureus isolates from the strain collection of the Andersson laboratory, Uppsala University, representing a range of infection sources: DA77312, DA77314, DA77316, DA77318, DA77320, DA77322, DA77324, DA77326, DA77328, DA77330, DA77332, DA81330, DA83662, DA83663, DA83664, DA83712, DA83713, DA83714, DA83715, DA83716, and DA83717. Isolate characteristics and their vancomycin and piperacillin–tazobactam minimum inhibitory concentrations are given in Extended Data Table 3.

### Growth media and culture conditions

All experiments were performed in Mueller–Hinton broth, with the exception of *E. faecium*, which was grown in lysogeny broth. Cultures were incubated at 37 °C with continuous shaking.

### Bacteriophage propagation and titration

Lysates were prepared from single plaques by plate lysis, as previously described ^13^. A single plaque was picked with a sterile stick, resuspended in 4 mL soft agar together with 0.1 mL of an overnight culture of the host, and poured onto a phage plate. After overnight incubation at 37 °C the soft agar was scraped and resuspended in 10 mL broth with approximately 0.5 mL chloroform, centrifuged, filtered through a 0.2 µm filter and stored at 4 °C. Titres were determined by soft-agar overlay on the corresponding host ^35^ and were re-checked before each experiment.

### Antibiotic panel and concentrations

All antibiotics were purchased from Sigma-Aldrich, and stock solutions were prepared fresh for each experiment. Every species received tetracycline (TET), chloramphenicol (CHL), piperacillin–tazobactam (TZP), and colistin (COL), allowing the same agent to be compared across bacterial backgrounds, together with two further agents selected for clinical relevance to that organism: vancomycin (VAN) and fosfomycin (FOS) for *E. faecium*; VAN and daptomycin (DAP) for *S. aureus*; ceftazidime (CAZ) and meropenem (MEM) for *K. pneumoniae* and for *A. baumannii*; ciprofloxacin (CIP) and CAZ for *P. aeruginosa*; MEM and trimethoprim– sulfamethoxazole (SXT) for *Enterobacter* sp.; and CIP and azithromycin (AZI) for *E. coli*. The two combination agents were used at fixed ratios: piperacillin–tazobactam at 8:1 and trimethoprim–sulfamethoxazole at 1:19 ^36^. Each antibiotic was used at half of its minimum inhibitory concentration for the species in question, as reported in Extended Data Table 2.

### Minimum inhibitory concentrations

Minimum inhibitory concentrations were determined by broth microdilution according to Clinical and Laboratory Standards Institute (CLSI) guidelines ^36^, at the inoculum specified in that standard, in three independent replicates, in the same medium in which each species was profiled in the growth-curve assay. The values are reported in Extended Data Table 2 for the seven-species panel and in Extended Data Table 3 for the *S. aureus* isolates.

### Growth-curve assay

Each combination was profiled in four conditions: untreated control, phage alone, antibiotic alone, and the combination. Experiments were performed in 100-well honeycomb plates in a Bioscreen C automated growth analyzer at 37 °C with continuous shaking, in a volume of 250 µL per well. Each well was inoculated with 10^7^ bacteria, and, in the phage and combination conditions, with 10^7^ phage, giving a multiplicity of infection of 1; overnight densities were estimated and phage lysates re-titred before each experiment so that this ratio was met. Antibiotics were applied at half the MIC, a concentration that produces measurable but incomplete growth inhibition ^9, 12, 14^ and leaves a growing population on which the phage can act. Optical density was recorded at 600 nm every 5 min for 48 h, with five independent replicates per condition ^23, 32^. This design parallels a time–kill experiment ^21^ but is miniaturized and parallelized, and it requires no instrumentation beyond the growth analyzer.

### Interaction metric

Raw optical density trajectories were transformed into relative growth curves by normalizing each condition to the untreated control at each time point, as previously established for growth-based interaction analysis ^23^. This normalization accounts for baseline growth variability and enables direct comparison of treatment effects across conditions and across organisms with different growth capacities.

For each experiment, the expected effect of independent action between phage and antibiotic was computed at every time point using a Bliss-based formulation ^22, 23^, in which the theoretical response is defined as the product of the relative growth observed for each agent alone. This generates a time-resolved reference trajectory representing non-interacting behavior. The experimentally observed growth under combined treatment was then compared with this theoretical expectation across the full duration of the assay.

Rather than relying on a single time point or growth phase, interaction strength was quantified by integrating both the experimental and the theoretical trajectories over time to obtain their respective areas under the curve (AUC). The difference between these quantities, ΔAUC = AUC(experimental) − AUC(theoretical), is the value of the sublethal interaction factor (SIF) for that combination ^23^. Deviations from zero indicate non-independent behavior: negative values correspond to enhanced inhibition relative to expectation (synergy), whereas positive values indicate reduced inhibition (antagonism), consistent with the original SIF formulation.

This framework yields a single quantitative endpoint per phage–antibiotic condition while preserving the full temporal information during analysis. Interaction effects arising at different phases of bacterial growth, including delayed lysis or transient recovery, therefore contribute to the final classification rather than being discarded.

Normalization to the untreated control assumes that all wells start from comparable densities. Differences present at time zero, arising from inoculation or blanking, propagate through the relative-growth curve and are compounded by the multiplicative Bliss expectation; in a small number of isolates such offsets were observed before any treatment effect could have occurred, and these coincided with the largest and least reproducible interaction values. An explicit quality-control criterion on time-zero agreement, assessed against a reference strain and phage, is recommended.

### Time-resolved interaction profiling

For the rolling formulation, the AUC was accumulated from the start of the experiment to each time point for both the observed combination and its theoretical expectation, and the difference between the two cumulative values was expressed as a percentage of the expected value at that time point: ΔAUC (%) = 100 × (experimental − expected) / expected. A value of +100 % therefore means twice as much growth as expected and −50 % half as much. Normalizing to the expected AUC makes species and isolates with different growth capacities directly comparable. The first hour is omitted throughout because the accumulated AUC is still near zero there and the ratio is unstable.

### Interaction classification

Because the absolute magnitude of ΔAUC scales with the growth capacity of the organism, a neutral interval around zero is treated as indistinguishable from independent action: |ΔAUC| < 5,000 for endpoint values and ±15 % for normalized values. Interaction calls are not made on deviations within that interval, so that assay noise is not reported as synergy or antagonism. The neutral interval used here was defined empirically and should be calibrated against replicate variance in each laboratory.

### Figure construction

Endpoint ΔAUC values are displayed on a symmetric logarithmic axis that is linear between −5,000 and +5,000 and logarithmic beyond it, so that both small and large interactions remain visible (Figs. 2 and 4). In Fig. 3a the y-axis is truncated at ±45 % and the four combinations exceeding it are marked with a triangle at the time of their maximum deviation, with that value given. In Fig. 2a, the four antibiotics tested in all seven species are shown left of the divider and the remaining antibiotics, tested in selected species only, to the right; blank cells were not tested. Row order in Fig. 5b, d is given by hierarchical clustering of each isolate’s joint VAN and TZP trajectory (Ward linkage on Euclidean distances, optimal leaf ordering ^37^), and the same order is used in both panels.

Antibiotics are abbreviated throughout as tetracycline (TET), chloramphenicol (CHL), piperacillin–tazobactam (TZP), colistin (COL), vancomycin (VAN), daptomycin (DAP), fosfomycin (FOS), ciprofloxacin (CIP), ceftazidime (CAZ), meropenem (MEM), azithromycin (AZI), and trimethoprim–sulfamethoxazole (SXT).

### Statistical analyses

All conditions were profiled in five independent replicates. Sample size was not predetermined by a power calculation. The identities of the bacterial isolates and of the antibiotics were randomized before the experiments, and B.A.B. blinded the data before analysis by N.F.K. No data were excluded. Each combination was tested against the null hypothesis of no interaction using a one-sided Wilcoxon signed-rank test ^38^ in the direction of the observed effect; with n = 5 the smallest attainable p value for this test is 0.031, so a single significance threshold (p < 0.05) is reported throughout. Error bars in Fig. 2b are the standard error of the mean across replicates; Fig. 4b shows the median and interquartile range across isolates.

Replicate variance scaled with effect magnitude: the standard deviation exceeded the absolute mean in 8 of 42 isolate–antibiotic combinations, so large effects require additional replication to be classified with confidence.

### Software and data analysis

All analyses were performed with a custom analysis pipeline, which takes raw plate-reader output and returns the endpoint and rolling interaction values, the classifications, and the figures. The pipeline is available as a web tool at https://github.com/nikoskava/SPARK. The hierarchical clustering uses Ward linkage on Euclidean distances with optimal leaf ordering.

### Reporting summary

Further information on research design is available in the Nature Portfolio Reporting Summary linked to this article.

### Web implementation

The analysis is implemented as a web tool at https://github.com/nikoskava/SPARK, which accepts raw plate-reader output and returns the interaction scores without local installation. The exported optical density file is uploaded as written by the reader, and the wells of each experiment are assigned to the four conditions of the design: untreated control, phage alone, antibiotic alone, and the combination. The tool then performs the steps described above, normalizing each condition to the untreated control at every time point, constructing the Bliss expectation from the two single-agent curves, and integrating the observed and expected trajectories. It returns the area under the curve for each condition, the endpoint ΔAUC, and the rolling ΔAUC expressed as a percentage of expected. Uploaded data are not retained server-side.

## Data availability

All data are available in the accompanying Data File.

## Code availability

The web implementation of the analysis is licensed under the MIT license and available at https://github.com/nikoskava/SPARK

## Author contributions

Conceptualization: BAB, NFK

Methodology: BAB, TGG, NFK

Investigation: BAB, TGG, DHS, NFK

Visualization: NFK

Funding Acquisition: NFK, DIA

Project Administration: NFK

Writing– Initial Draft: BAB, DHS, NFK

Writing– Review & Editing: BAB, TGG, DHS, DIA, NFK

## Acknowledgements

We would like to thank Daria Van Tyne, Madison Stellfox, Breck Duerkop, Arya Garima, Patrick Kenney, Edze Westra, Anthony Maresso, Liu Pengbo, Marlene Wolfe, Christine Moe, and Orlando Sablon III for their assistance in acquiring suitable phages for this project. We would like to thank the U.S. National Institute of General Medical Sciences for their funding support via R35GM136407 to Bruce R. Levin and the Swedish Research Council for their funding via a VR grant 2017-01527 to Dan I. Andersson. The content is solely the responsibility of the authors and does not necessarily represent the official views of the National Institutes of Health or the Swedish Research Council

## Extended Data

**Extended Data Table 1.** Bacteria and phage identities and their sources.

| **Bacteria** | **Bacteriophage** | **Phage Source** |
|---|---|---|
| *Enterococcus faecium* | Φ4 | Daria Van Tyne |
| *Staphylococcus aureus* | PYO | Levin Lab |
| *Klebsiella pneumoniae* | LL1 | Isolated |
| *Acinetobacter baumannii* | Φ1P | Patrick Kenney |
| *Pseudomonas aeruginosa* | DMS3<sup>Vir</sup> | Edze Westra |
| *Enterobacter sp.* | Φ13 | Daria Van Tyne |
| *Escherichia coli* | ES19 | Anthony Maresso |

**Extended Data Table 2.**
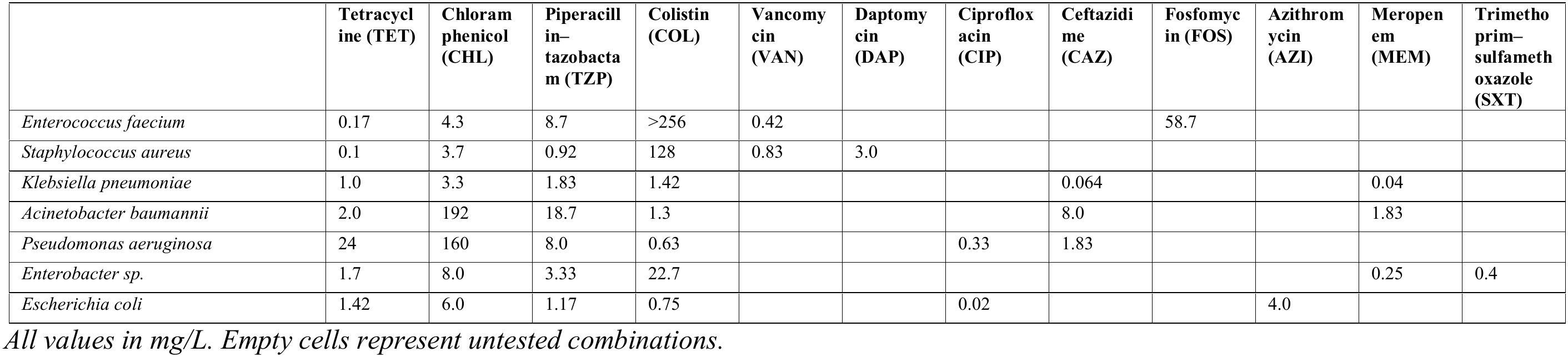
Minimum inhibitory concentrations of the twelve antibiotics against the seven species of the panel.

**Extended Data Table 3.**
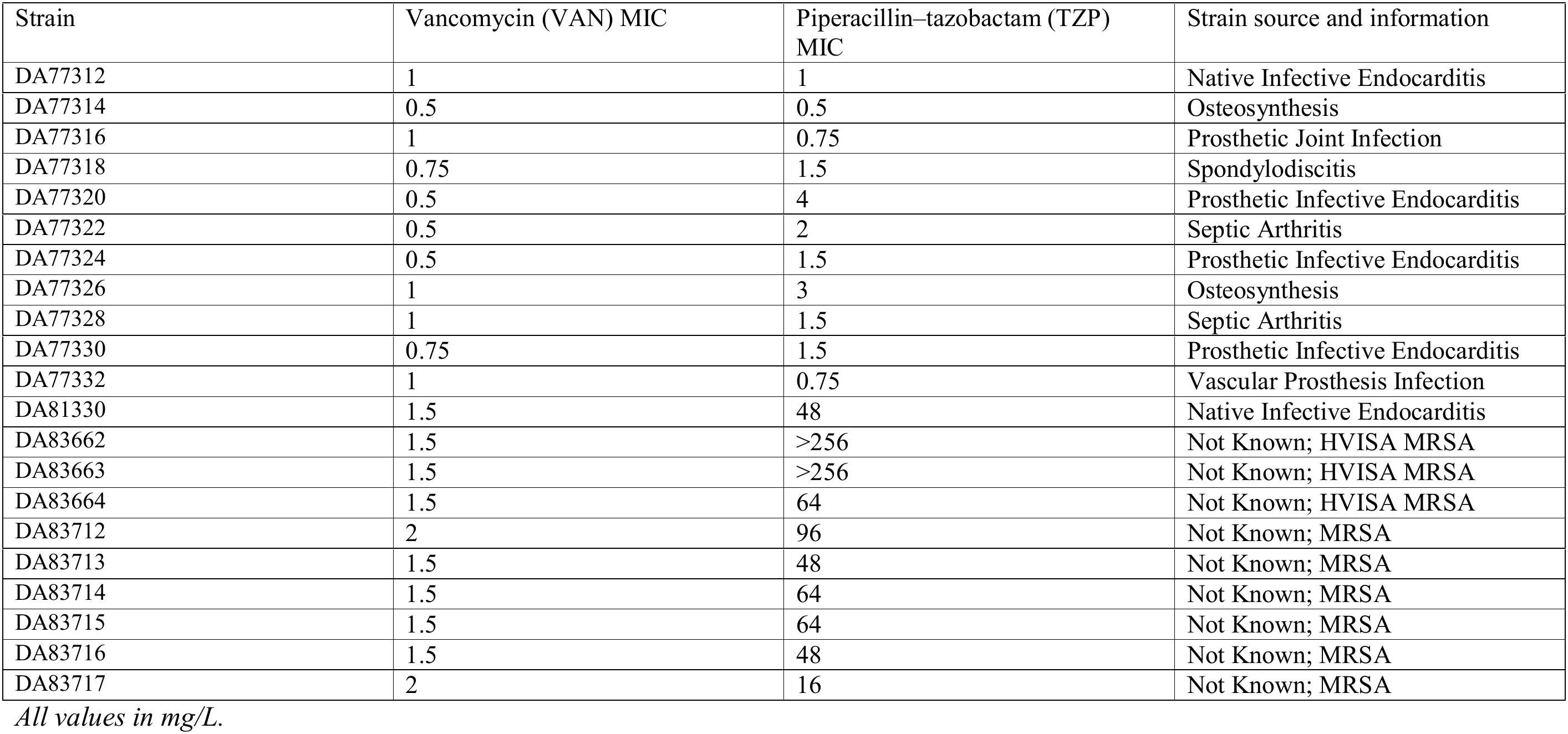
*Staphylococcus aureus* isolates, their minimum inhibitory concentrations, and isolate information.

